# A spatio-temporal individual-based network framework for West Nile virus in the USA: spreading pattern of West Nile virus

**DOI:** 10.1101/438366

**Authors:** Sifat A. Moon, Lee W. Cohnstaedt, D. Scott McVey, Caterina M. Scoglio

## Abstract

West Nile virus (WNV)—a mosquito-borne arbovirus— entered the USA through New York City in 1999 and spread to the contiguous USA within three years while transitioning from epidemic outbreaks to endemic transmission. The virus is transmitted by vector competent mosquitoes and maintained in the avian populations. WNV spatial distribution is mainly determined by the movement of residential and migratory avian populations. We developed an individual-level heterogeneous network framework across the USA with the goal of understanding the long-range spatial distribution of WNV. To this end, we proposed three distance dispersal kernels model: 1) exponential—short-range dispersal, 2) power-law—long-range dispersal in all directions, and 3) power-law biased by flyway direction—long-range dispersal only along established migratory routes. To select the appropriate dispersal kernel we used the human case data and adopted a model selection framework based on approximate Bayesian computation with sequential Monte Carlo sampling (ABC-SMC). From estimated parameters, we find that the power-law biased by flyway direction kernel is the best kernel to fit WNV human case data, supporting the hypothesis of long-range WNV transmission is mainly along the migratory bird flyways. Through extensive simulation from 2014 to 2016, we proposed and tested hypothetical mitigation strategies and found that mosquito population reduction in the infected states and neighboring states is potentially cost-effective.

**Author summary:** The underlying pattern of West Nile virus (WNV) geographic spread across the United States is not completely clear, which is a necessary step for continental or state level mitigation strategies to reduce WNV transmission. We report a network model that explains the geographic spread of WNV in the United States. West Nile virus is a mosquito-borne pathogen that infects many avian species with different movement ranges. From our research, we found that migration patterns and routes play an essential role in the WNV spatial distribution. The virus spreads in all directions at short distances because of local birds and short-distance migratory birds. However, the virus also disperses long distances along the avian migratory routes. Our model is designed to be flexible and therefore can be used to explore spreading patterns of other infectious diseases in other geographic locations.

## Introduction

West Nile disease (WND) is a vector-borne zoonosis which may result from infection by West Nile virus (WNV), a member of the family Flaviviridae, genus Flavivirus. This virus is the most common cause of arboviral disease in the United States [1]. From 1999 to 2017, more than 48 thousands WNV disease cases were reported to the Centers for Disease Control and Prevention (CDC) and more than two thousands of these reported cases resulted in death [2]. WNV is maintained in an enzootic transmission cycle between competent mosquitoes and birds. Birds are the reservoir and amplifying host for this virus. The US Centers for Diseases Control and Prevention (CDC) has identified WNV infection in more than three hundred species of birds. Infected bird movement is likely a key factor that affects the geographic spread of WNV, especially given the different habitats and routes of various species. Although many bird species may be infected with WNV, the American robin is considered an important amplifier of WNV and maybe a driver geographic spread because WNV-infected American robins have low mortality and high viremia [3, 4]. Members of the *Culex* genus of mosquito are the principal vectors of this virus in the United States [5]. Humans, horses, and other mammals can be infected with WNV. However, these infections result in relatively low virus titers (viremia) therefore the infected animals and people are considered dead-end hosts (not capable of infecting feeding mosquitoes). Therefore, they do not have any epidemiological impact on WNV transmission or geographic spread [6].

To understand the transmission dynamics of WNV, several mathematical models have been developed [3, 7–10]. These models predict the threshold conditions for WNV spreading in different scenarios. However, most of these models do not consider the spatial dynamics of WNV. Space or geographic spread has a significant role in WNV disease dynamics and modeling of WNV spatial spreading is complex because of the interactions of multiple potential mosquito vectors, avian amplifiers, and mammalian hosts. Liu et al. [9] developed a patchy model to analyze the spatial spreading of WNV, where patches are geographical space. They assumed patches are identical, spatial dispersal of birds and mosquitoes are symmetric within patches, and movement of birds and mosquitoes are only one-dimensional. According to this investigation, long-range dispersal of infected bird populations determines the spatial spread of WNV, not the dispersal of infected mosquito populations. Other investigators proposed a reaction-diffusion model [10], where they have spatially extended the non-spatial model of Wonham et al. [8] to mathematically estimate the spread of WNV. Here, diffusion terms in the reaction-diffusion partial differential equations represent vector mosquito and host bird population movements. They identified traveling wave solutions in their model and calculated the rate of spatial spread of infection. Durand et al. [11] developed a discrete time deterministic meta-population model in order to analyze the circulation of WNV between Southern Europe and West Africa. Another spatial model proposed by Maidana and Yang [12] used a system of partial differential reaction-diffusion equations. They also calculated the speed of disease dissemination by investigating the traveling wave solution of their model. They concluded, mosquito movements do not play an important role in disease dissemination. In addition, they included vertical transmission in their model and determined that vertical transmission is not an important factor for the spatial spread of WNV.

Most WNV spread models are mathematical deterministic compartmental models. However WNV spread is highly stochastic because of the demography and movement of hosts and vectors varies between different locations. The major weaknesses of these models are the number and complexity of the compartments required to account for the many host and vector populations. In turn, the number of compartments increases the number of unknown parameters. Approximation of these parameters in any biological system is very challenging and prone to estimation errors which can create inaccuracies in the model outputs.

We developed an individual-based heterogeneous network framework to understand WNV geographic spread. To build the network framework, we used the American Robin population density across the contiguous United States. The demographic characteristics of avian host populations and vector populations are not homogenous geographically, so we used a heterogeneous network framework. The transmission intensity of WNV depends on the abundance of WNV-infected vector mosquitoes in a given location. Mosquito population numbers fluctuate with local weather and season throughout the year, therefore we used a temperature dependent transmission rate. Although dead-end hosts cannot spread WNV to mosquitoes, we have quantified WNV case data only for humans, which we used to estimate unknown parameters.

To understand the WNV spatial distribution, we proposed distance dispersal kernels, which describes the probability of dispersal with respect to distances. In this framework, we proposed three types of distance dispersal kernels: 1) exponential, 2) power-law, and 3) power-law biased by flyway. Then we compared the three distance kernels using approximate Bayesian computation based on sequential Monte Carlo sampling (ABC-SMC) method [13–18]. After conducting an extensive simulation for 2014-2016, we observed that an adapted fat-tailed or power-law kernel, which has long-distance links in specified directions can best describe the WNV human case data. We tested this network framework for the best kernel with the human case data and found that simulated results for more than 41 states of 49 states are consistent with the reported WNV cases. We proposed several theoretical mitigation strategies to control WNV and calculated their estimated costs. From the analysis of mitigation strategies, we suggest that potentially effective mitigation policies would include the application of mitigation control in areas with active transmission and in immediate neighboring states.

## Materials and methods

In this section, we present data sources, an epidemic model for WNV, then develop a network framework for WNV geographic spread across the United States. At the end of this section, we present a statistical tool, approximate Bayesian computation with sequential Monte Carlo sampling (ABC-SMC) for parameter estimation and model selection.

### Data

The study area of this research was the contiguous United States where WNV is considered endemic. We modeled WNV case distributions for 2014-2016. We used three data sets each year to develop our model. The first dataset contained the average monthly temperatures. Mosquito vector abundance correlated with temperature. Temperature data was from the National Centers for Environmental Information [19]. The second dataset contains American Robin population data from *eBird* [20]. This is a database for bird abundance and distribution, which is formed by the Cornell Lab of Ornithology and National Audubon Society. We used total observation of American Robin in each state of the USA for each month. The robin data set was used to train the network model. The American Robin is abundant throughout the United States and is a preferred food source for many WNV-competent mosquito species [21]. Based on host feeding patterns of the *Culex* genus of mosquitoes, robins are the most common WNV amplifying host [22–24]. Other important susceptible birds, such as American crow were not used because although they are an indicator species (high crow mortality), they are unlikely to spread virus geographically as they are mostly a residential species. In addition, as an indication of epidemic start point, we used WNV human incidence data. Many species of birds have long-distance migration during the spring and fall. Therefore the network does not focus on one long-distance migrating bird species but aggregates all species along the known flyways. To estimate model parameters we used human case data for WNV from CDC [2], which is the third dataset.

### WNV Epidemic model

To explore WNV long-distance spatial distribution in the USA, we used an individual-based heterogeneous network framework. In this framework, birds are on the individual level, a node represents an individual bird and connection between nodes is the possibility of virus dispersal from one infected bird to another susceptible bird by mosquito vectors. Links or connections are formed by movement of birds or movement of vectors. If there is no link between nodes then infected birds and insects are not moving virus between nodes. All virus transmission occurs by local competent vector mosquitoes. There is some evidence of bird-to-bird transmission, but it likely does not contribute to or maintain outbreaks. We split the bird population into four compartments; susceptible, exposed, infected, and recovered. Although, in the literature most mathematical models do not consider the exposed avian class when modeling WNV [8, 12, 25, 26]. Birds transmit virus to mosquitoes when a susceptible mosquito vector takes an infected blood meal, then the mosquito becomes infectious after the extrinsic incubation period (EIP), or the time needed for the virus to spreads from the mosquito mid gut to the salivary glands; usually this process takes 7 to 14 days [3, 27]. In addition, an infected bird can infect many mosquitoes simultaneously and also an infected mosquito can bite many susceptible or infected birds. Therefore, there is some delay in the system, to represent this delay we added the exposed class. We estimated exposed period from data by using the approximate Bayesian computation with sequential Monte Carlo sampling (ABC-SMC) method. After the exposed period, birds entered the infected compartment and an infected bird transitions to recovered after 4-5 days. To simulate this model, we used generalized epidemic mean-field (GEMF) framework developed by the Network Science and Engineering (NetSE) group at Kansas State University [28]. In GEMF, each node stays in a different state and the joint state of all nodes follows a Markov process [28–30]. The node level description of this Markov process is:

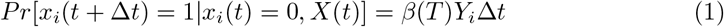

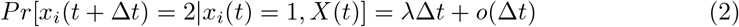

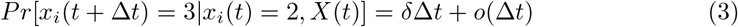

Here, *X*(*t*) is the joint state of all individual nodes at time *t*. *x_i_*(*t*) is a node state, *x_i_*(*t*) = *C* means node *i* is in *C* compartment at time *t*, *C* = 0, 1, 2, 3 corresponds to susceptible, exposed, infected, and recovered compartment. *Y_i_* is the number of infected neighbors of node *i*, *β*(*T*) is the probability of transmission from one infected bird to one susceptible bird, which is a function of temperature, *λ* is the rate for exposed to infectious state, and finally, a node recovers from infectious state at a rate *δ*.

#### Zoonotic spillover transmission

To model disease transmission from the bird population to human population, we added a zoonotic spillover transmission compartment. We modeled occurrence of human cases as a Poisson process [26, 31]. This part of the framework can be expressed as the following equation:

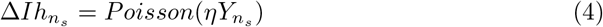

In this equation, *Ih_n_s__* is number of infected human cases at *n* sub-network in *s* time steps, where *s* = 1, 2, 3….. are the discrete time steps, *Y_n_s__* is infected bird population in sub-network *n*, and *η* is a scaler quantity, accounts for the contact rate and probability of pathogen transmission from bird to human. We calculated WNV spilling over to humans by using a Poisson random number generator.

#### Temporal transmission rate and environmental conditions

The transmission rate for WNV is sensitive to weather data as mosquito abundance depends on the environmental conditions. Temperature, precipitation, landscape features, daylight conditions etc. are environmental conditions, which has an impact on the transmission dynamics of WNV [32]. In this research, we considered average monthly temperature data, optimal mosquito season [33], and suitable temperature range for co-occurrence of WNV and competent mosquito species. Temperature plays a very important role in the transmission dynamics of WNV because mosquito longevity and EIP are sensitive to temperature. Mosquito longevity and EIP decrease with the increase of temperature. However, there is no straightforward relationship of vectorial capacity for WNV with temperature. If incubation period decreases more than longevity, then mosquitos will be infective longer. However if longevity decreases more than incubation period, then mosquitos will not be able to transmit the virus. We used information about rainfall in this research implicitly through optical mosquito season. Optimal mosquito season of any location is estimated from monthly average temperature and rainfall data for that location [33]. In this model, we used a simple linear relation of transmission rate with temperature in a temperature window from 12°C to 32°C in the optimal mosquito season. Outside this window, transmission rate is very low. Suitable temperature for co-occurrence of WNV and *Culex pipiens* is around 12° to 27°C and for *Culex quinquefasciatus* is 20°C to 32°C [33]. Survival rate to adult stage for *Culex quinquefasciatus* is significantly high when temperature is in 20°C to 30°C [34]. For *Culex tarsalis* favorable temperature for WNV development start after 14°C [35], however larval survival reduced after 30°C temperature [36]. To compute the transmission rate of any link from node *a* to node *b*, we used temperature of the location of node *b*. Transmission rate for a location *l* is, *β_l_*(*T*) = *β_○_*(*T_lm_* − *T_○_*); here, *β_○_* is the proportional constant, what we estimated by using ABC-SMC method, *T_lm_* is the average temperature for month *m* in location *l* and *T_○_* is the threshold temperature. Threshold temperature for this model is 12°C. As the temperature is space dependent, our transmission rate also differs across the network. This individual-level heterogeneous network model gives us this flexibility to use different transmission rate at a time for different parts of the network.

### Network framework

For the spatial dynamic characteristics of WNV transmission, we built a network framework, which has 49 sub-networks one for each adjoining states of the contiguous United States plus the District of Columbia. The number of nodes in each sub-network is proportional to the size of the avian population in that state [20]. We considered the mosquito season June-October for the simulation period. Although the mosquito season is not the same for all states, mosquitoes are active from June to September in all of the states at these times [33].

The network for the avian population is (V, E). Here, *V* is the set of nodes, which is the union of nodes of all sub-network, *V* = *SN* 1 ∪ *SN* 2 ∪ *SN* 3 ∪ ……….. ∪ *SN* 49, here *SNi* is a set of nodes in the sub-network *i* and *E* is the set of links among individual nodes. To build sub-networks, we used the total number of observations of American Robin for states per month in the simulation time period. 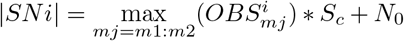, here, 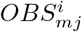 is the total number of observations of American Robins in state *i* in month *mj*, *N*_0_ is the error term and *N*_0_ ~ *N* (5, 2) for this model. *m*1 is the first month after May and *m*2 is the last month before October when the average monthly temperature is greater than *T*_0_. *S_c_* is the scaling constant.

In each sub-network, we assumed that nodes are connected through *Erdos-Renyi (n,p)* random network topology [37]. In this network topology, we created links randomly among nodes with a probability *p*. Here, n is the number of nodes in a sub-network and *p* is the probability to form an edge. We set the probability *p = R * log*(*n*)*/n*, here *R* is a constant (*R* ≥ 2), as this value is more than the threshold value for the connectedness of an *Erdos-Renyi* graph [38], so nodes of a sub-network are locally connected. We will refer these networks as a local network in the subsequent sections of this paper. To build connections among sub-networks, we considered long-distance dispersal kernels, which describe the probability of dispersal with respect to distances. Dispersal kernels provide a simple model of dispersal to model dispersal events. For long-distance events, we used three types of kernel models; 1) Exponential, 2) power-law, and 3) power-law-flyway, which is a power-law kernel biased by flyway. A simple caricature of the network is shown in Fig 1. There are three sub-networks, A, B, and C. The links, which formed local networks are shown by solid lines. These links are introduced by *Erdos-Renyi (n,p)* network topology. Dashed lines are inter-links among sub-networks. These links established by using long-distance dispersal kernels.

**Fig 1.**
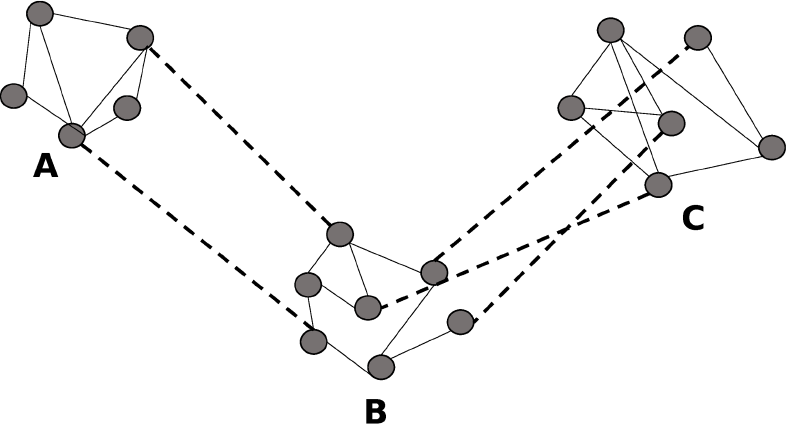
A simple caricature of the actual contact network for the avian population. Here, A, B, C are three sub-networks. Solid lines represent intra-links in a sub-network and dashed lines represent inter-sub-network links.

#### Exponential distance kernel

In this distance kernel, connection probability among sub-networks will decrease exponentially with distance. Probability to form a link is:

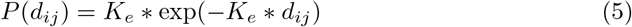

Here, *d_ij_* is the distance between sub-network *i* and *j*, *K_e_* is the shape parameter of exponential distribution kernel. For distance between two states, we took the distance between their centroids. The network with the exponential dispersal kernel was created as follows:

Step 1 Calculate the distance among sub-networks. *d_ij_* is the distance between sub-network *i* and *j*.
Step 2 Calculate *P* (*d_ij_*), this is the probability to form a link between sub-network *i* and *j*.
Step 3 Generate a random number *rand* for each pair of nodes *(a,b)*, where *a* ∈ *i* and *b* ∈ *j*.
Step 4 If *rand* < *P* (*d_ij_*) then an undirected link will form between node *a* and *b*.

Inter-links among sub-networks, generated by exponential distance kernel are shown in Fig 2a.

**Fig 2.**
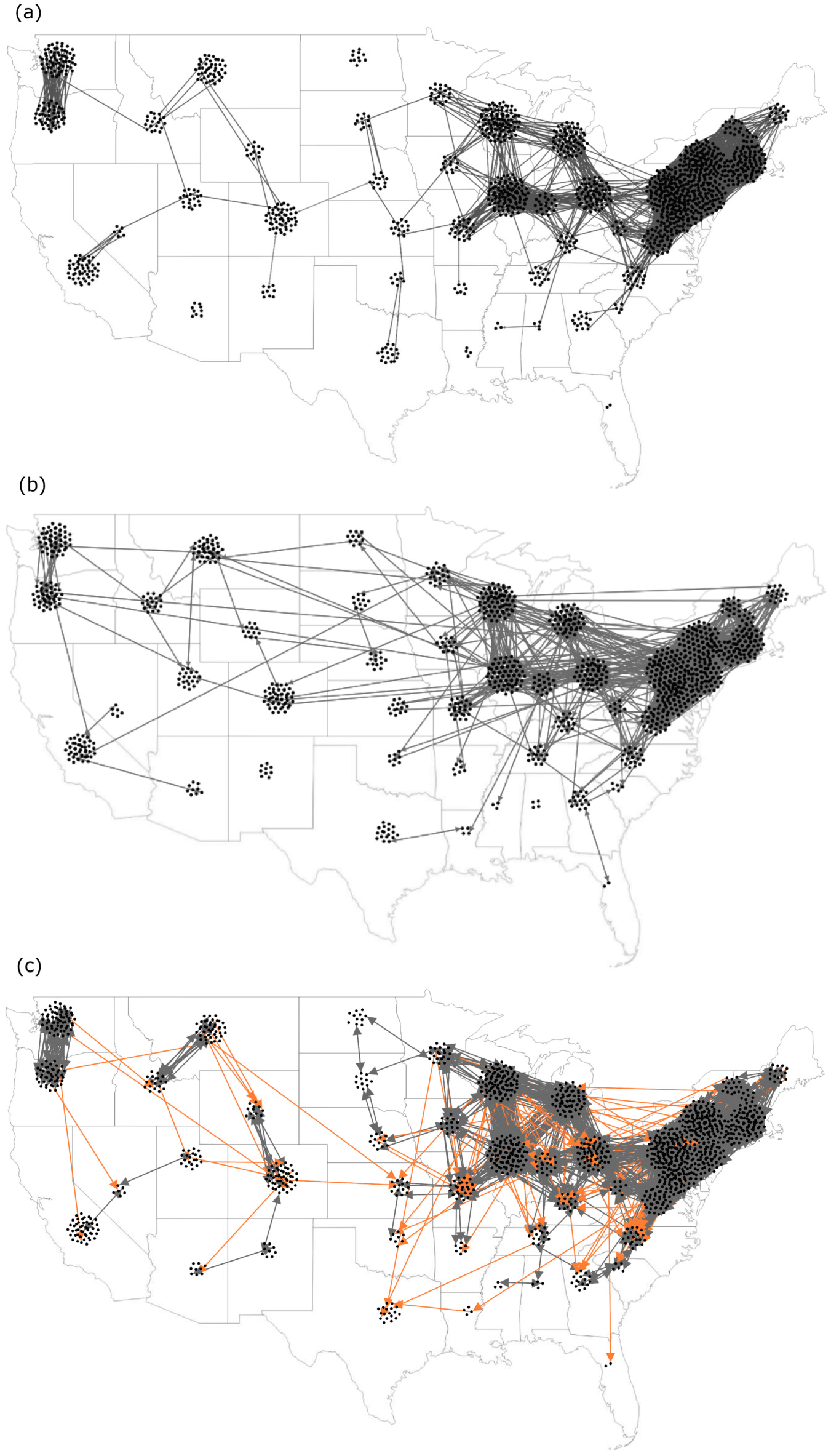
Inter-links among sub-networks. a) for exponential distance kernel, b) for power-law distance kernel, and c) for power-law distance kernel biased by flyway. Gray links represent undirected links and orange links represent directed links (for spring migration–northbound; for late summer/fall migration–southbound). Intra-links are not visible here. These are one realization of the stochastic networks, which are rescaled by 0.1 for better visualization.

#### Power-law distance kernel

Power-Law, heavy-tailed, or fat-tailed distribution allows occasional long-range transmissions of infection with frequent short-range transmissions. In this fat-tailed distance kernel, there is a greater chance of creating links over the same long-distances compared to the exponential kernel. Power-law transmission kernel was used previously to model spatial dynamics of several infectious diseases, for example, in plant epidemiology [39], in 2001 foot-and-mouth disease epidemic [40], and also, in human diseases [41]. In power-law connections [42], the probability of connectivity among sub-networks will decrease with distance according to the following equation:

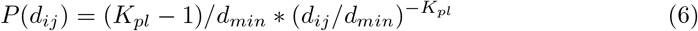

Here *d_min_* is minimum distance among sub-networks and *K_pl_* is the power-law parameter. The process to build this network is similar to a network for exponential kernel with the only difference being the calculation of *P* (*d_ij_*). Inter-links among sub-networks for power-law distance kernel are shown in Fig 2b.

#### Power-law distance kernel biased by flyway

To form this distance kernel, we included the migratory behavior of birds. Migratory birds can spread pathogens during the migration periods [43, 44]. According to the United States Fish and Wildlife Services and Flyway Councils, there are four flyways in the United States; the Atlantic flyway (AF), the Mississippi flyway (MF), the Central flyway (CF), and the Pacific flyway (PF) [45]. Although flyways overlap and the migratory patterns are very complex, these migratory routes play a vital role in the long-distance spreading of WNV [46]. To build this distance kernel, we considered two types of links among sub-networks; 1) links which are formed for residential or short-distance migratory bird movements and 2) links which are formed for long-distance migratory bird movements. For the first type of links, we used an estimated movement range of 500 km [47], these connections are unrelated to flyways. For the second type of connections, we considered two migration periods; spring migration (April - June) and late summer/fall migration (July - September) [30]; during the spring migration, we established long links from south to north and in late summer/fall migration, the reverse. To establish any long link, we picked two sub-network and establish a link if they were in the same flyway with probability *P* (*d_ij_*) (Eq. 6), these links were directional and direction was imposed with respect to migratory period. Inter-links among sub-networks for this kernel were shown in Fig 2c. The algorithm to create this network was:

Step 1 Calculate the distance among sub-networks. *d_ij_* is the distance between sub-network *i* and *j*.
Step 2 Calculate *P* (*d_ij_*) using Eq. 6, this is the probability to form a link between states *i* and *j*.
Step 3 Generate a random number *rand* for each pair of nodes *(a,b)*, where *a* ∈ *i* and *b* ∈ *j*.
Step 4 If *rand* < *P* (*d_ij_*) and *d_ij_* < 500*km* then an undirected link will form between node *a* and *b*.
Step 5 If *rand* < *P* (*d_ij_*) and *d_ij_* > 500*km* and states *i* and *j* are in the same flyway then an directed link will form between node *a* and *b* according to the migration period.

#### Temporal network behavior

Bird populations are not constant in any region, they change with time because of bird movement. To consider this fact, this study adds a node property, namely, *Activity*. This property can hold two values: 1 = *Active* and 0 = *Inactive*. In the entire network, only *Active* node can contribute to the spreading of the WNV. By controlling this property, we varied the size of the active node population in any sub-network with respect to the variation of the avian population in that region. The length of the simulation each year was five months (June - October). Then, each month nodes are activated randomly according to the total number of birds observed in that region in that month.

### ABC-SMC for parameter estimation and model comparison

In this framework, we adopted approximate Bayesian computation based on a sequential Monte Carlo sampling (ABC-SMC) method for parameter estimation and model selection [13–18].

#### Parameter estimation

ABC-SMC is a computational method of Bayesian statistics that combines a particle filtering method with summary statistics. This method is ideal for a stochastic complex model where likelihood function is intractable or computationally expensive to evaluate. ABC estimates the posterior distribution of parameters from data. Let, *θ* is a parameter vector to be estimated. The goal of the ABC is to approximate the posterior distribution, ∏(*θ|d*) ∝ *f* (*d|θ*)∏(*θ*), where prior distribution of parameters ∏(*θ*) are given and *f* (*d|θ*) is the likelihood of *θ* given the data *d*. This method samples parameter values from their prior distribution through subsequent SMC rounds. Intermediate distribution of the parameter is ∏(*θ|dist*(*x*, *d*) ≤ *ε_i_*); *i* = 1, 2,…. *P*. The target posterior distribution is ∏(*θ|dist*(*x*, *d*) ≤ *ε_P_*). Here, *x* is the simulated data set, *dist* is the distance function, *ε* is the tolerance and *P* is the number of SMC rounds or the number of populations, where *ε_P_* < ….. < *ε*_2_ < *ε*_1_ [48]. This is an adapted sequential importance sampling. In each SMC round, it uses perturbation kernel to sample a parameter set. After each simulation of the model, the model output and data are compared using some goodness-of-fit metrics. A parameter set is accepted if the distance between the model output and data is less than the tolerance level. The accepted parameter set is a particle and accepted particles form a population for that SMC round. We used two goodness-of-fit metric or distance function in this research. The first goodness-of-fit metric is squared root of the sum of squared error between observed incidence data and simulated incidence data for any proposed parameter set. The first goodness-of-fit metric for this model is:

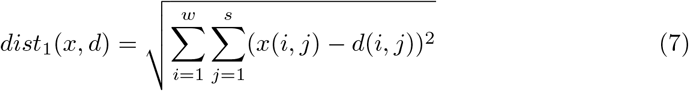

Here, *x(i,j)* is simulated incidence model data for *i* week and for *j* location. The second goodness-of-fit metric is the absolute difference between the number of infected states from observed data and simulated data, infected state defined as a state where at least one infected individual has reported. The ABC-SMC algorithm, we adopted for this model from Toni et al. [13], which has given in Text S2. We used this algorithm separately for estimating parameters for this three distance dispersal kernel network models. As our models are an event based stochastic simulation, we simulated them 30 times with GEMF for each particle to get 30 realizations of the system. Then we take the average of these realizations. As the average over the multiple runs of a stochastic system holds more information than a single stochastic run.

#### Model comparison

In many areas, researchers deal with model selection. Bayesian theory is a comprehensive method to make inference about models from data. Approximate Bayesian computation was used in many research areas for model selection [49]. To compare among three distance kernels, this investigation used ABC-SMC model selection framework [13, 50, 51]. For given data *d*, the marginal posterior probability of model m is:

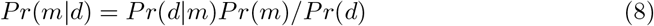

Here, *Pr*(*d|m*) is the marginal likelihood and *Pr*(*m*) is the prior probability of the model. We used a uniform distribution for prior distribution of unknown parameters. For each model, we have four unknown parameters; network parameter *K* (*K_e_* is the network parameter for the exponential kernel and *K_pl_* is the network parameter for the both power-law kernels), constant for transmission rate *β*_0_, transition rate from exposed to infectious state *λ*, and zoonotic transmission spillover rate *η*. In each population, we took 1000 particles. We used Bayes factor to compare a model with another model. For model *m_i_* and *m_j_*, Bayes factor [52] is,

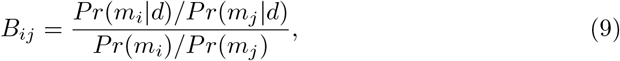

Here, *Pr*(*m_i_*) is the prior and *Pr*(*m_i_|d*) is the marginal posterior distribution of model *m_i_*. The Bayes factor is a summary evidence in favor of one model over another supported by the data. If *B_ij_* is in range 1-3, we can conclude that summary of the evidence against *mj* in favor of *m_i_* is very weak. If *B_ij_* is in range 3-20, we can conclude that summary of the evidence against *m_j_* in favor of *m_i_* is positive [52]. The ABC-SMC model selection algorithm is very similar to the algorithm for parameter estimation. Here, *m* is the model indicator, *m* ∈ 1, 2, …‥, *M*, *M* is the number of model. In this research, we had three network models (*M* = 3) to compare.

m = 1: exponential kernel network model,

m = 2: power-law kernel network model, and

m = 3: power-law kernel influenced by flyway network model.

In each population, the model selection algorithm starts by sampling the model parameter *m* from the prior distribution ∏(*m*). Then the algorithm proposes a new set of parameters (particle) from the sets of parameters of the model *m* from the previous population. The Bayes factor was calculated from the final population of *m*. The algorithm for model selection has given in Text S2. Although ABC-SMC is an accurate statistical tool for parameter estimation and model selection, however, the results of this method are sensitive to summary statistics [53]. For our case, no summary statistics were required because we used the entire set of data and we compared the simulated and observed dataset directly by using goodness-of-fit or distance metric. A full dataset is sufficient to get the consistent result from approximate Bayesian Computation [54].

### Mitigation strategies

The role of mosquito populations in WNV transmission is expressed by disease transmission rate *β*. This framework used different transmission rates in different parts of the network corresponding to the local mosquito abundance. Using this heterogeneous feature in the framework, we evaluated theoretical mosquito population management measures to reduce the outbreak size or transmission rates in the state level. Some states such as Kansas, do not have statewide mosquito surveillance or management, but in these theoretical scenarios, it is assumed they can develop or benefit from effective statewide mosquito management programs. The framework will simply estimate how much the mosquito abundance is reduced or maintained based on the theoretical outcomes of coordinated control. Furthermore, we realize mosquito control is generally conducted on a county or municipal level, but the human case data is only available on a state level. Therefore the recommendations are for the lowest resolution of the data, which is state level but applies to counties and municipalities as well. If vector management is increased in a sub-network, then transmission rates will be changed by, 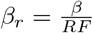, here *β_r_* is the reduced transmission rate and *RF* is the reduction factor. Then management costs will be *Cost* = *RF* * *NS*_*c*_, here *NS_c_* is the number of states where control measures were applied. We considered supplemental management measures with the existing management measures. We used two types of mitigation strategies across the United States, 1) dynamic infected place tracing strategy and 2) static ranked based strategy.

In the infected place tracing, we traced the infected states, then plan the mitigation strategies according to them. For this type of mitigation strategies, we considered three cases; 1) *case-1: only infected*: applied control only in the infected states; 2) *case-2: infected & first neighbors*: applied control in the infected states with its first neighboring states (whose distance is less than 500*km*), and 3) *case-3: infected & first neighbors & second neighbors*: applied control in the infected states with its first neighboring states, and also with its second neighboring states (whose distance is in 500 – 1000*km*). For infected tracing control measure, we kept track of infected places monthly. If *SNi* sub-network is infected for month *t*, then control measures were applied for the month *t* + 1 based on these three cases.

In the static ranked based mitigation strategy, we ranked the states by different variables (for example, temperature, size of the avian population etc.). For this strategy, we considered three cases; 1) *temp*.: states ranked by temperature, 2) *pop*.: states ranked by avian population size, and 3) *temp. & pop*.: states ranked by temperature and avian population size both, then we applied management measures in the top 30% of the states.

## Results

We developed a novel flexible individual based heterogeneous network framework to test three WNV dispersal kernels across the contiguous United States based on human case data distributions. We used this framework for the year 2014, 2015, and 1016. The results for network formulation, parameter estimation, and dispersal kernels selection using Bayesian inference are given below for the year 2015 and the results for other two years are given in the Tex S1.

### Network framework

In this spatial-temporal individual-based heterogeneous network framework, we used three distance kernel models. The fundamental basic WNV epidemic model is the same for all the three network kernels. In the entire network, there are 49 sub-networks representing the 48 adjoining contiguous states plus the District of Columbia. All sub-network nodes are locally connected. The topology of the local network is *Erdos-Renyi*. The total nodes for the year 2015 was *|V|* = 7657 and the scaling constant is *S_c_* = 0.02. Here, *E* = *E_l_* ∪ *E_dd_*; *|E_l_|* is the number of total intra-links for all local networks, which is around 167000-170000 and *E_dd_* is the number of total inter-links among sub-networks. The description of sub-networks is provided in S4 Table in the Text S3. We started the epidemic from states with the highest human incidence prior to June. We started the epidemic for the year 2015 by adding two infected nodes, one in sub-network *SN4* (California) and another in sub-network *SN42* (Texas). Connections among sub-networks are developed by distance dispersal kernels. Parameters for these kernels are estimated from the ABC-SMC method.

### ABC-SMC for parameter estimation and model comparison

#### Parameter estimation

ABC-SMC parameter estimation was applied to three dispersal kernel network models separately. For each set of prior distributions, convergence to the posterior distribution was achieved after 13-15 SMC rounds. Convergence of the posterior distributions was monitored by visual inspection of the outputs from consecutive SMC rounds. The prior distribution for exponential network parameter was, *K_e_ ~ U* (0.1, 0.3), for power-law *K_pl_ ~ U* (2, 4), for power-law biased by flyway was *K_pl_ ~ U* (2, 4). Prior distribution for constant of transmission rate *β*_0_, transition rate from exposed to infectious *λ*, and human spillover rate *η* is same for three kernel models; *β*_0_ *~ U* (0, 15), *λ ~ U* (0.025, 10) and *η ~ U* (0, 50). Perturbation kernels were also uniform, *PK* = *αU* (−1, 1), with *α* = 0.5(*maxθ*_*p*−1_ − *minθ*_*p*−1_), here *θ*_*p*−1_ is the set of a parameter values in the previous population. We used weekly human case data for 49 locations, as observed data. The estimated parameters for this three dispersal kernel network models for 2015 are presented in Table 1.

**Table 1.**
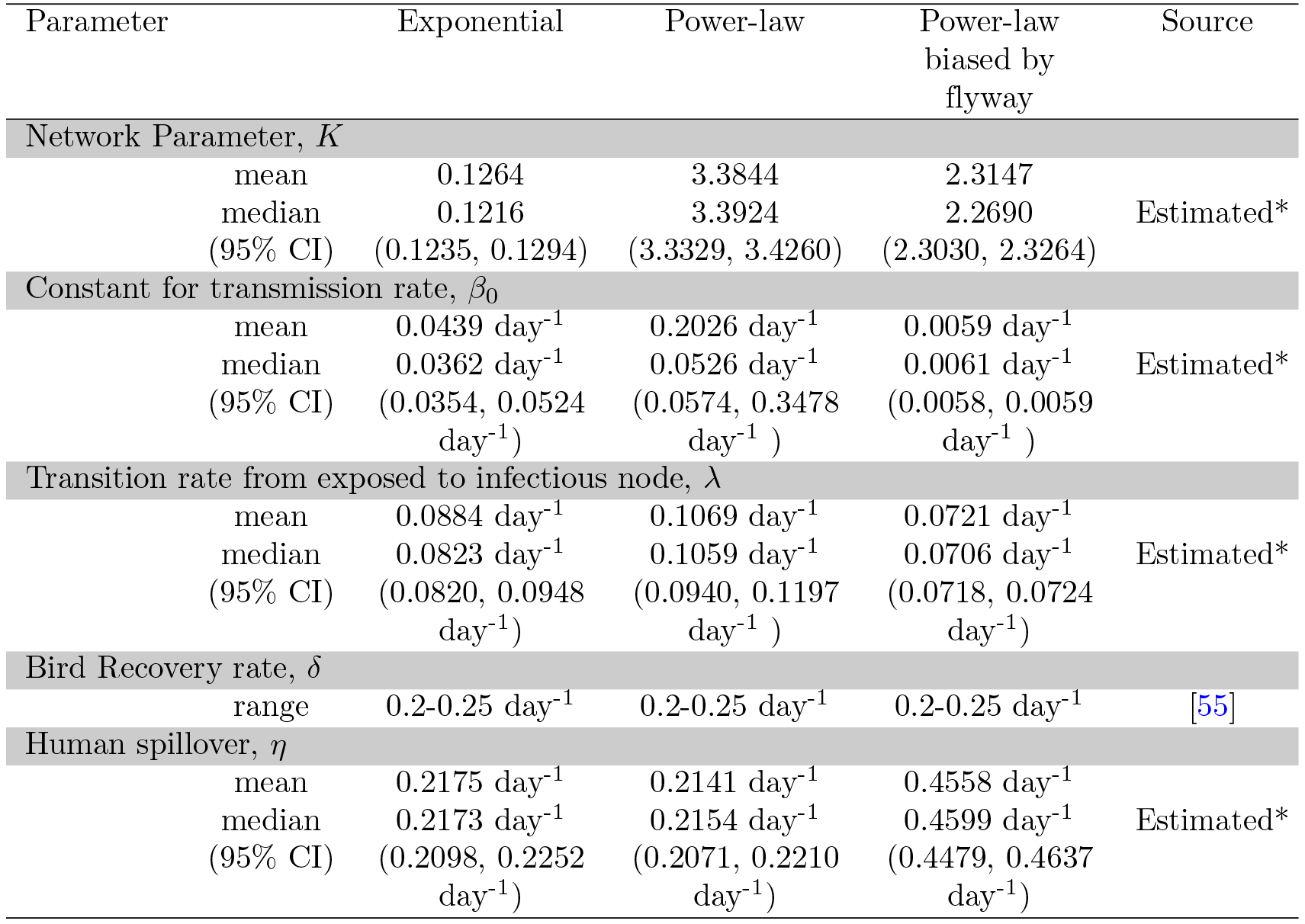
Estimated parameters for the year 2015 from ABC-SMC parameter estimation. *Estimated using data from the Centers for Disease Control and Prevention (CDC) [2], the National Centers for Environmental Information [19], and Clements et al. [20].

#### Model comparison

ABC-SMC for model selection allows us to estimate posterior model distributions. We used this algorithm to compare the three distance kernels. Prior distributions and perturbation kernels are the same for both the model selection and the parameter estimation algorithm. Here we used one more prior distribution for discrete model parameter; *m ~ U* (1, 3). The tolerance vector for ABC-SMC model selection algorithm is, *ε* = 2200, 2000, 1800, 1600, 1400, 1200, 1100, 1000. The target and intermediate distributions of model parameters are shown in Fig 3.

**Fig 3.**
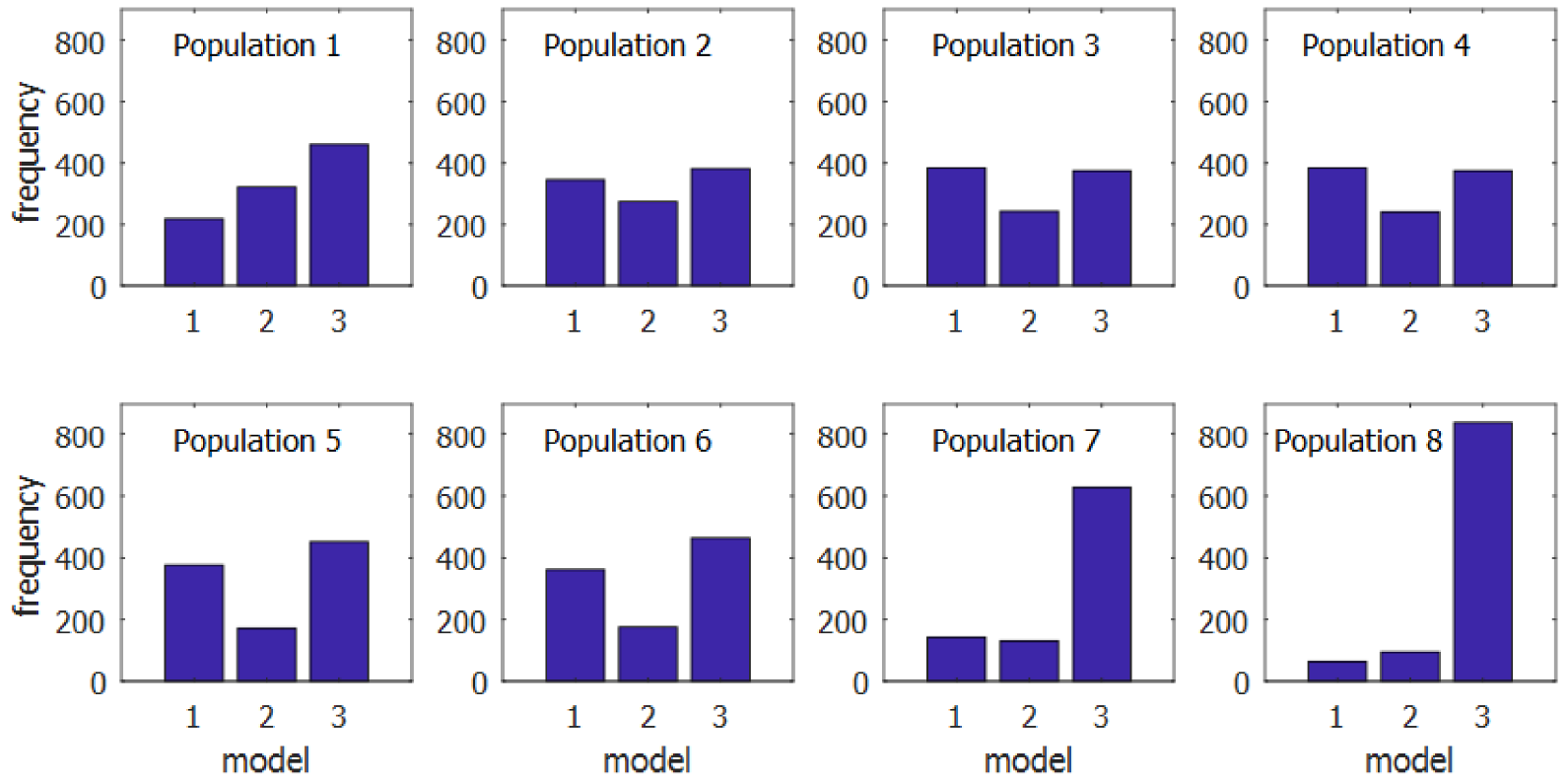
Population of the marginal posterior distribution of the three models for the year 2015. Model-1 represents exponential kernel, model-2 represents power-law kernel, and model-3 represents power-law influenced by flyway kernel. Here, Population-8 is the approximation of the final marginal posterior distribution of model parameter *m* and population 1-7 are intermediate distributions. Population-0 is the discrete uniform prior distribution, which is not shown here.

We calculated the Bayes factor from the marginal posterior distribution of *m*, which we took from the final or last population. In the final population for 2015, exponential distance kernel model (*m* = 1) was selected for 64 times, power-law distance kernel (*m* = 2) was selected for 95 times and power-law influenced by flyway distance kernel model (*m* = 3) was selected for 841 times. Bayes factor *B*_3,1_ = 841/64 = 13.1406, *B*_3,2_ = 841/95 = 8.8526. In the marginal posterior distribution of three models, there is positive evidence in favor of power-law influenced by flyway distance kernel when compared with other two models [13]. The distribution of parameters for power-law influenced by flyway for 2015 are presented in Fig 4. Calculation of the Bayes factor for 2014 and 2016 are provided in the Tex S1.

**Fig 4.**
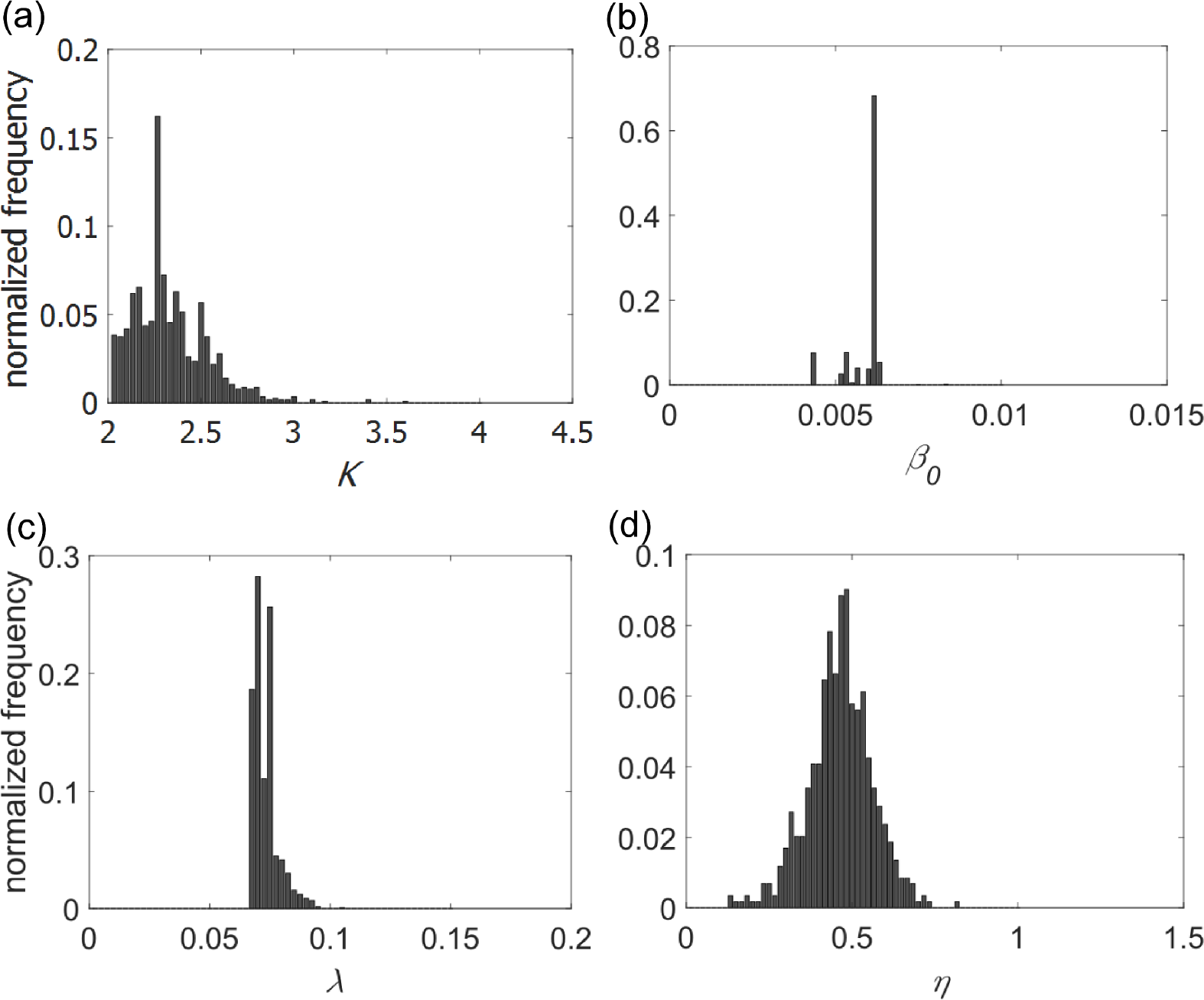
Histograms of the approximated posteriors distribution of parameters for power-law influenced by flyway kernel for the year 2015. a) Network Parameter *K*; b) constant for transmission rate *β*_0_; c) transition rate from exposed to infectious node *λ*, and d) human spillover *η*.

### Performance of the power-law-flyway network model

To test the performance of this framework, we used estimated parameters from Table 1 for power-law kernel influenced by flyway. We set the parameters value; *K_pl_* = 2.3147, *β*_0_ = 0.0059*day*^−1^, *λ* = 0.0721*day*^−1^, and *δ* = 0.2031*day*^−1^. The simulation period for the avian population model is from week-23 to week-44. The output of avian population was used as the input of zoonotic spillover compartment. Then we compared the output of zoonotic spillover compartment with human case data for week 24 to week 45. We considered a one-week lag between WNV incidence in birds and WNV incidence in humans. In humans, WNV-infected individuals (approximately 20%) develop a mild febrile illness after 3–6 days [56]. Peak of reporting of dead birds is one week prior than the reporting peak of human incidence [57]. In Fig 5, the mean simulated human case from the 49 sub-networks is compared with the weekly human case data for 2015 for the contiguous USA. The absolute errors between them are shown here. From this whisker plot, we can see that the median of the absolute error for the states is close to zero. In Fig 5, the largest outlier is California (marked by black circles). These outliers result from a mismatch between the simulated peak human incidence time and the observed human incidence peak time possibly because the very long state (north to south) has weather which is very different in southern California (warmer and drier) than northern California (cooler and wetter) causing a difference between peak mosquito seasons in the southern and northern parts.

**Fig 5.**
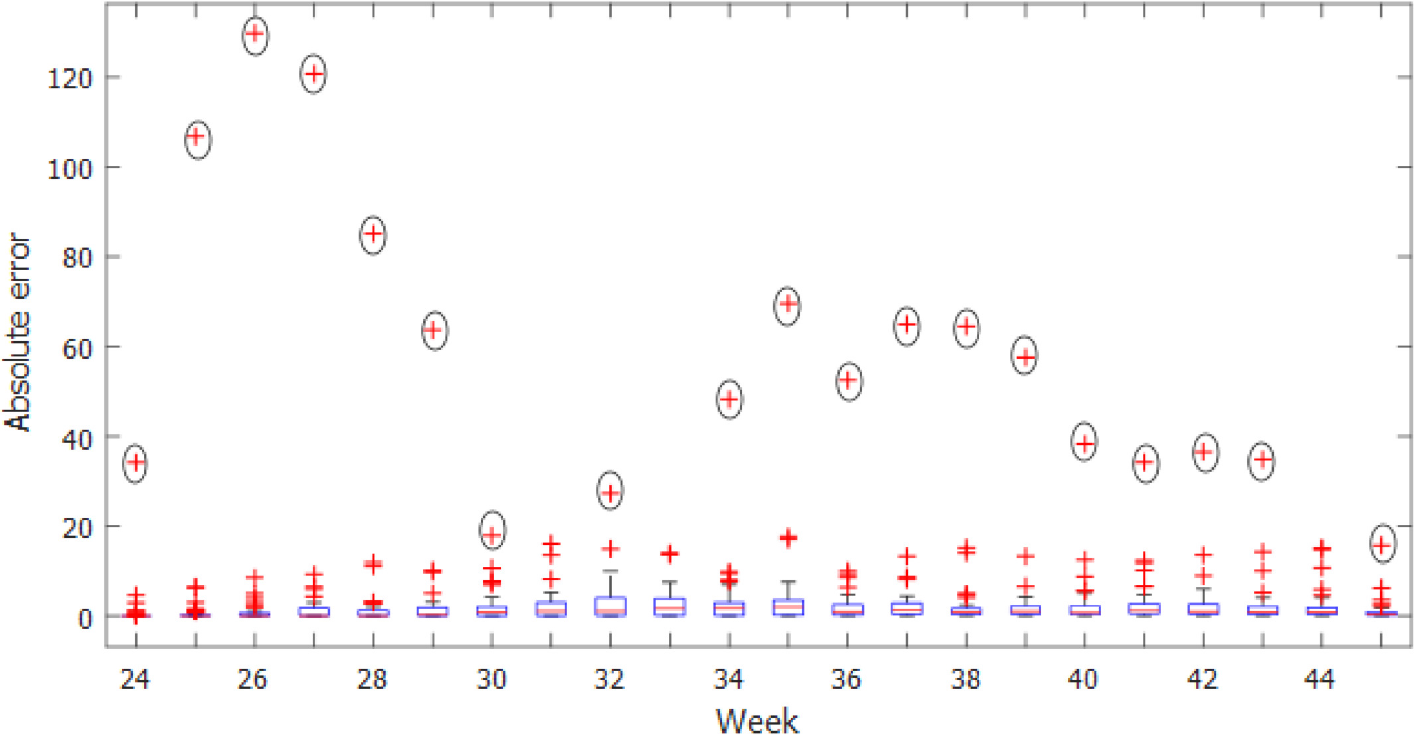
Absolute errors of the simulated human cases of 49 states by weeks with the observed data for the year 2015. Mean of 1000 realizations has used as the simulated data. On the blue boxes, the red horizontal lines show the median and the bottom and top edges of the boxes indicate 25^th^ and 75^th^ percentile respectively. The whiskers show the ranges of data points not considered outliers and outliers are showing by red + symbol. Californian outliers are marked by black circles.

We compared the total yearly incidence of human WNV from this model with the state level reported case data. The results are shown in Fig 6. For 2015, we found that the case data for 42 of 49 locations were within the simulation results. The states where human cases were different from the simulation results were *over-reported* states (Nevada) and *under-reported* states (Louisiana, Mississippi, Nebraska, North Dakota, South Dakota, and Washington). The possible reason for this mismatch are reporting error or overwintering of virus in birds or mosquitoes or another bird species (not robins) is the key reservoir species for that state

**Fig 6.**
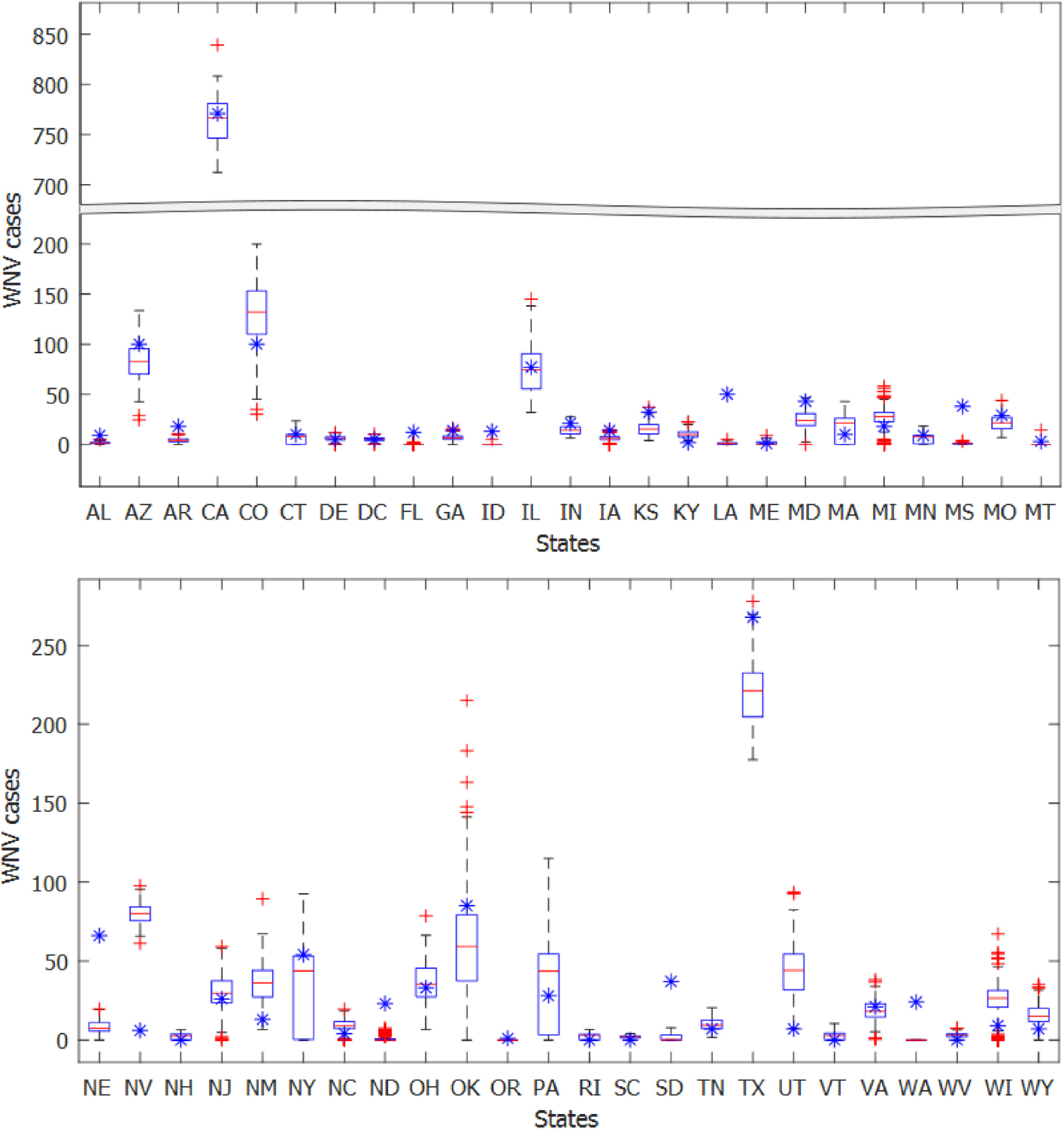
WNV human incidence by states for the year 2015 from power-law influenced by flyway kernel model. (for *K_pl_*=2.3147, *β*_0_ = 0.0059*day*^−1^, *λ* = 0.0721*day*^−1^, *η* = 0.4558*day*^−1^), generated from 1000 simulation and observed data are indicated by blue colored star points. states name are given in the short form. Simulated results are represented with a box plot in which the red horizontal lines show the median and the bottom and top edges of the boxes indicate 25^th^ and 75^th^ percentile respectively, The whiskers show the ranges of data points not considered outliers and outliers are showing by red + symbol. Broken scale is used for sake of visualization.

To build a disease prevalence map, we grouped the states in four categories; 1) higher prevalence—incidence is more than 100, 2) intermediate prevalence—incidence is in between 50-99, 3) moderate prevalence—incidence is in between 25-49 and 4) low prevalence—incidence is less than 25. To group the states, we used the median of the simulation results. The disease prevalence map from the model are presented in Fig 7a and from observed data are presented in Fig 7b. Among 49 locations, 40 locations are in the same prevalence group in both maps.

**Fig 7.**
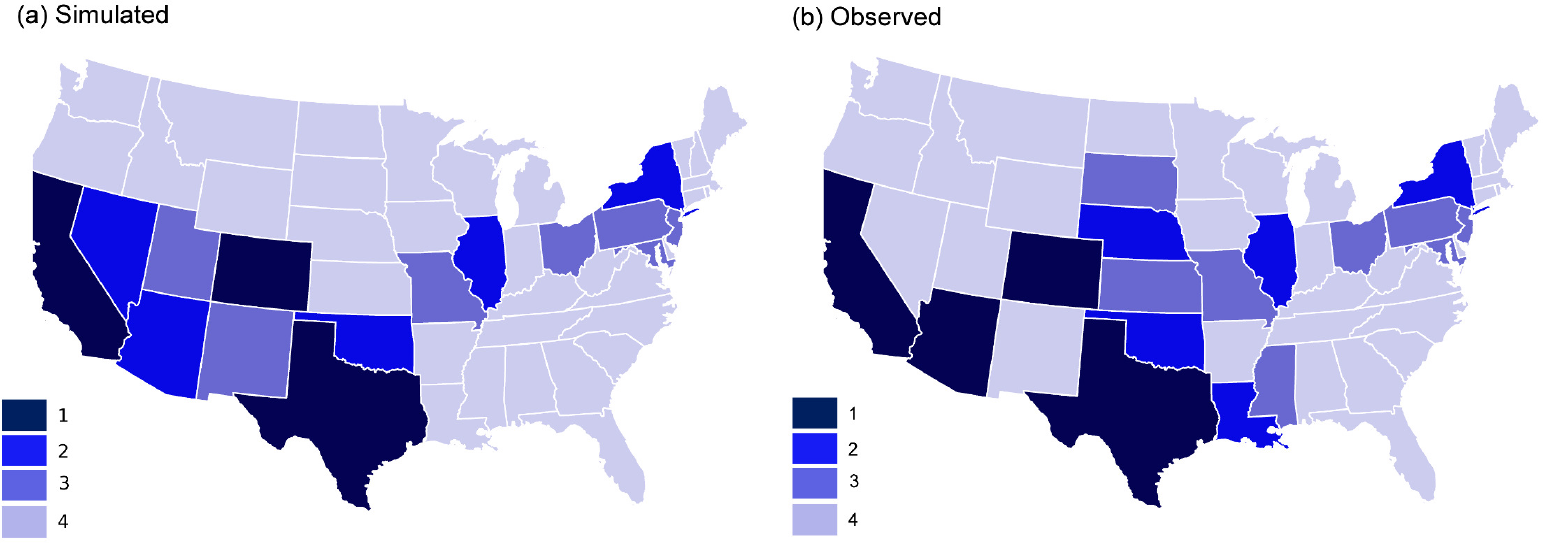
Disease prevalence map for WNV human incidence for the year 2015. The darker regions imposed greater prevalence. States are divided into four groups by incidence number; group-1: more than 99, group-2: 50-99. group-3: 25-49, and group-4: less than 25 incidences. a) States are divided by the median of the output of 1000 simulations, b) states are divided by observed data.

### Mitigation strategies

We applied mitigation strategies on the power-law-flyway kernel network model to find the optimal mitigation plan. Fig 8a shows the number of infected states or epidemic size for dynamic infected places tracing. Epidemic size decreased faster with increased reduction factor for *case-2* (infected & first neighbors) and *case-3* (infected & first neighbors & second neighbors) than *case-1* (only infected). The number of states where control measures were applied is displayed in Fig 9, which is proportional to cost. Therefore, the cost was minimal for case-2 than other two cases for *RF* > 2. From the cost analysis, we concluded that, although the cost for *case-1* is less at the beginning of the yearly outbreak, we need to apply management only in the infected places, however by the end of the year the total cost for *case-2* will smaller because of the smaller epidemic size.

**Fig 8.**
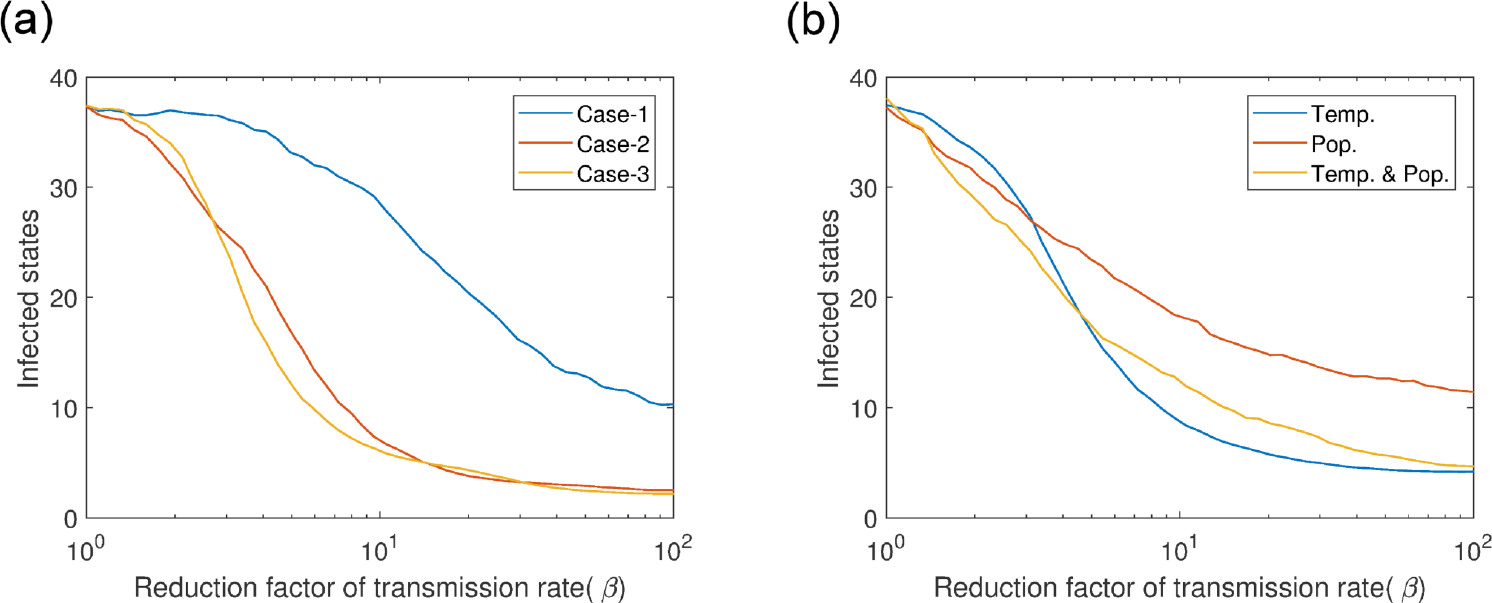
Infected states for two types of mitigation strategies. a) Dynamic infected places tracing; case-1: control measures are applied only in the infected states, case-2: control measures are applied in the infected states plus in their first neighboring states, case-3: control measures are applied in the infected places plus in their first and second neighboring states, and b) static ranked based strategy–states are ranked by; 1) temperature (Temp.), 2) avian population size (Pop.), 3) both(Temp & Pop.), then control measures are applied in the top 30% states. Log scale has used in x-axis for better visualization.

**Fig 9.**
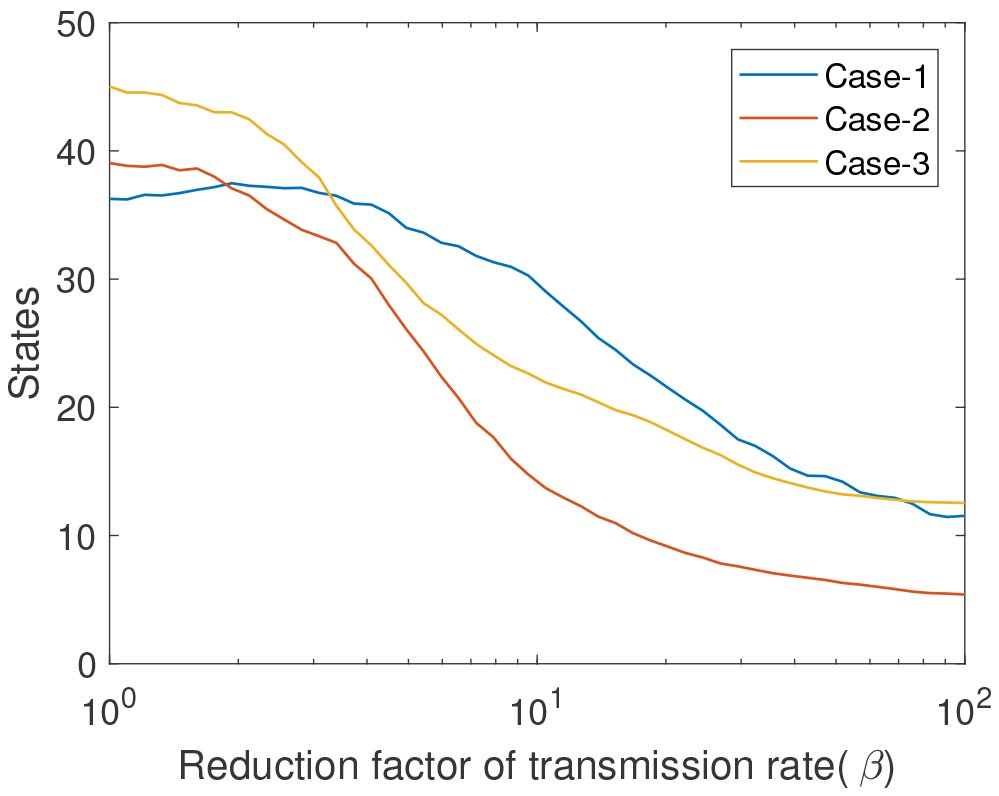
Number of states where control measures are applied for the infected places tracing mitigation strategy. Log scale has used in x-axis for better visualization.

The results of the static ranked based mitigation strategy measure are presented in Fig 8b. We observed that, before *RF* = 4.5, number of infected states for *temp. & pop*. dropped earlier than others. Number of infected states or epidemic size was smaller for *temp*. than *pop*. after *RF* > 3, infected population of a sub-network are more positively correlated with temperature. The *NS_c_* is always the same for these three cases. For all mitigation strategies, minimum epidemic size could be 2, as we started the epidemic from two states.

## Discussion

We proposed an individual-based heterogeneous network framework and tested three dispersal kernels to understand the spatial spread patterns of WNV human case data across the contiguous United States.

This framework requires fewer parameters and has more flexibility to represent the spatial-temporal dynamics of WNV. Adding parameters will make the framework more realistic, for example, more competent bird species, landscape features for habitat preferences of host and vector species, daylight conditions [32], pathogen invasion from outside of USA, variable susceptibility among different hosts and vectors, WNV strain variability, mosquito and virus overwintering, vertical transmission, human movement characteristics etc․. However, inclusion of too many factors increases model complexity which makes model optimization difficult given the availability of limited observational data. On the other hand, a simple model may insufficient to represent WNV spatial dynamics. Computational models need to be developed and parameters calculated with sufficient detail to be biologically accurate if they are used to evaluate epidemic management measures. However, for most biological systems, reliable parameter information is unknown. Unknown parameters or inaccurate assumptions add uncertainty to the model. Our framework has only four parameters to estimate (network Parameter *K*, transmission rate *β*, transition rate from exposed to infectious state, *λ*, and human spillover, *η*). This framework has compartments only for the avian population (susceptible, exposed, infected, and recovered), which does not have to be species specific. We reduced the compartments for vector population by implementing them implicitly through transmission rate between infected nodes and susceptible nodes. The presented framework and dispersal kernel network model has an intermediate complexity that approximate Bayesian computation based on sequential Monte Carlo sampling (ABC-SMC) method successfully calibrated and estimated the parameters with the available data. If more data becomes available, it is possible to add them in this model for improved performance of the model.

Furthermore, this framework is flexible and therefore can represent various hosts and vectors including with population seasonality, which plays an important role in WNV dynamics. For host population seasonality, we added a node property *Activity*, this property allows us to control active host populations in the network in a specific time period. We added vector seasonality in this framework through temperature dependent transmission rate. This framework proposed one exponential and two fat-tailed distance kernel models for long-distance transmission of WNV. WNV spatial distribution is very complex because WNV can infect more than 300 bird species, some of which are residential birds and short-distance migrators which disperse less than 500 km distances (short connections) whereas some species are long-distance migratory birds creating long connections. The long-distance migratory birds are the long-distance dispersal (LDD) agents for WNV. Previous studies tried to analyze spreading of WNV using a traveling wave with constant velocity, however, WNV spread more rapidly across the North America than would be expected from the assumption of constant velocity traveling wave [58]. Likely this is because traveling wave models unlike distance dispersal kernel models for WNV spreading do not capture the long-distance migrating birds which can have various migratory ranges and distances. Distance dispersal kernels have more flexibility to represent the different bird migration distances and can account for accelerating invasions. However, exponential kernels produce short-connections and therefore like traveling waves are limited to constant expansion, unlike fat-tailed power-law kernels which can generate accelerating invasions by creating the long-distance connections from migratory birds [59]. However, a general fat-tailed power-law kernel makes long-distance links in every direction which does not follow the incidence of WNV. Instead, a power-law-flyway kernel can be used to produce the long connections in the direction of flyways and short links in other directions. Bayesian inference was used to test which of the three kernel models best described WNV distribution on the network for three most recent years (2014- 2016). The power-law-flyway kernel best described the distribution of WNV cases because the long-range WNV transmission was concentrated mainly along the migratory bird flyways. The general power-law kernel overestimated the incidence data in some states because it was creating long-distance links in all directions.

The performance for the power-law-flyway dispersal kernel model was evaluated for the three most recent years (2014-2016) when WNV was endemic in the USA. The observed case data for the 49 locations were within the range of the simulated results for 41 states for 2014 (Fig S2 in Tex S1), 42 states for 2015 (Fig 6), and 45 states for 2016 (Fig S4 in Tex S1). For all three years, the simulated results were similar to the observed data, except in Colorado, Louisiana, Mississippi, Nevada, Nebraska, North Dakota, and Washington. Nevada was *over-reported* for 2015 and all others were *under-reported*. The power law flyway dispersal kernel network model reported more WNV human incidence in Nevada than reported cases, one possible reason for over-reporting cases in Nevada has rural areas, which tend to under report human cases, whereas mosquito control districts and health departments, focused in urban areas, must test birds and mosquitoes, which explains why CDC reported WNV infected mosquitoes in 25% of counties in Nevada. The under-reported states had more human cases than predicted by the model. Under-reporting by the power-law-flyway kernel network model is likely because overwintering of the virus in some states (for example, Louisiana, Mississippi etc.), which was not considered. The overwintering infected *Culex* mosquitoes can stay in hibernacula such as sewers, houses, caves, and other warm areas in urban, suburban, and rural areas and initiate the outbreak in the spring. Furthermore, there may be under-reporting of cases by the model if robins are not the main reservoir species in a state, which would be predicted between gulf coast states (Louisiana and Mississippi) and northern states such as North and South Dakota and Washington.

Mitigation strategies for WNV were tested using the power-law dispersal kernel network model. The management measures are not specific to larvae or adults, rather simply generally accepted best practices to reduce mosquito abundance for the purpose of reducing pathogen transmission. The mitigation strategy analysis proposes supplemental measures in addition to the existing mosquito management in each state because the states had yearly reported WNV cases despite the existing management methods. To reduce WNV spread, a theoretical policy would be management in neighboring regions and not exclusively in the infected places. Although this approach can cost more at the beginning of the epidemic season however at the end, it can reduce total cost by decreasing the size of the epidemic. If management measures are applied only in the infected states, it is not possible to control the epidemic because of long-distance migratory birds. This is a statewide management in a unified effort. We acknowledge that states do not conduct mosquito management in this way, but to test the spillover it was necessary to do the simulation in this way because only state level data was available.

Cooperation and communication equal early treatment and reduced outbreak sizes because of reduced WNV dispersal by American robins. This novel model can be applied to find out the invasion patterns of other long-distance dispersing pathogens.

## Supporting information

**Tex S1. Supplementary Materials.** Simulation results for 2014 and 2016.

**Text S2. Approximate Bayesian Computation based on sequential Monte Carlo sampling (ABC-SMC) method**.

**Text S3. Network description for 2014-2016**.

## Acknowledgements

We would like to thank Dr. Christopher Mundt for his excellent guidance in our EEID (Ecology and Evolution of Infectious Diseases) project (USDA: 2015-67013-23818, 3020-32000-008-04-S). We also would like to thank the developers of the GEMF software [28].

